# Gene length and detection bias in single cell RNA sequencing protocols

**DOI:** 10.1101/119222

**Authors:** Belinda Phipson, Luke Zappia, Alicia Oshlack

## Abstract

Single cell RNA sequencing (scRNA-seq) has rapidly gained popularity for profiling transcriptomes of hundreds to thousands of single cells. This technology has led to the discovery of novel cell types and revealed insights into the development of complex tissues. However, many technical challenges need to be overcome during data generation. Due to minute amounts of starting material, samples undergo extensive amplification, increasing technical variability. A solution for mitigating amplification biases is to include Unique Molecular Identifiers (UMIs), which tag individual molecules. Transcript abundances are then estimated from the number of unique UMIs aligning to a specific gene and PCR duplicates resulting in copies of the UMI are not included in expression estimates. Here we investigate the effect of gene length bias in scRNA-Seq across a variety of datasets differing in terms of capture technology, library preparation, cell types and species. We find that scRNA-seq datasets that have been sequenced using a full-length transcript protocol exhibit gene length bias akin to bulk RNA-seq data. Specifically, shorter genes tend to have lower counts and a higher rate of dropout. In contrast, protocols that include UMIs do not exhibit gene length bias, and have a mostly uniform rate of dropout across genes of varying length. Across four different scRNA-Seq datasets profiling mouse embryonic stem cells (mESCs), we found the subset of genes that are only detected in the UMI datasets tended to be shorter, while the subset of genes detected only in the full-length datasets tended to be longer. We briefly discuss the role of these genes in the context of differential expression testing and GO analysis. In addition, despite clear differences between UMI and full-length transcript data, we illustrate that full-length and UMI data can be combined to reveal underlying biology influencing expression of mESCs.

## Background

Single cell RNA-Seq (scRNA-Seq) has rapidly gained popularity as the primary tool to profile gene expression of hundreds to hundreds of thousands of single cells. This new technology enables researchers to examine transcription at the resolution of a single cell in a high-throughput manner, and has led to the discovery of novel cell types and revealed insights into the development of complex tissues as well as differentiation lineages. With the promise of novel discoveries, this new technology has been embraced by the scientific community.

Many technical challenges need to be overcome during data generation, and technology for performing scRNA-Seq is advancing at a rapid rate. The original Fluidigm C1 system has a 96 well plate which limits how many single cells researchers can practically handle in an experiment. However, depth of sequencing is only limited by cost, with sequencing depth of around 2 million reads per cell recommended (Tung et al. 2016). Droplet based technology, such as InDrop (Klein et al. 2015), Drop-Seq (Macosko et al. 2015) and the more recent Chromium system from 10X Genomics (Zheng et al. 2016), are cost effective methods to obtain relatively shallow sequencing of thousands to tens of thousands of single cells in one run. The lower sequencing depth per cell limits the complexity of the expression profile attained as only the most highly expressed genes will be observed, however, it may be the case that researchers combine deeper sequencing of fewer single cells with shallow sequencing of tens of thousands of cells to answer their scientific questions of interest.

Not only are there different technologies for capturing single cells, there are also differences in library preparation protocols, which aim to amplify and process the minute amounts of RNA from each cell. Most RNA-Seq library preparation protocols include enrichment of mRNA by either polyA pulldown or ribosomal depletion, followed by fragmentation and PCR amplification before sequencing. The extensive PCR amplification that is required for scRNA-Seq increases technical variability in the data by introducing amplification biases (Stegle et al. 2015). A solution for mitigating amplification biases is to include Unique Molecular Identifiers (UMIs), which are short (5-10bp) sequences ligated onto the 5’ end of the molecule prior to PCR amplification (Islam et al. 2013). Transcript abundances are then estimated from the number of reads with unique UMIs aligning to a specific gene. PCR duplicates resulting in copies of the UMI are therefore not included in expression estimates. While some protocols, such as those used with Fluidigm C1 (e.g SMARTer), need to be modified to include UMIs, some droplet based methods, for example, the Chromium system, always include UMIs in the chemistry. It is worth noting that while mechanisms such as alternative splicing can be studied using full-length transcript protocols, this type of analysis is not possible with data generated with protocols that include UMIs.

Gene length bias is well understood in bulk RNA-seq data. When cDNAs are fragmented, long genes result in more fragments for the same number of transcripts, resulting in higher counts and more power to detect differential expression (Oshlack & Wakefield 2009). As a result gene set testing is biased towards gene ontology categories containing longer genes (Young et al. 2010). While there is much in common between scRNA-Seq and bulk RNA-Seq data, modifications to the protocols such as amplification and the inclusion of UMIs, may highlight different biases in the data.

Here we investigate the effect of gene length bias in scRNA-Seq across a variety of datasets differing in terms of capture technology, library preparation, cell types and species. As hypothesised, we find that scRNA-seq datasets that have been sequenced using a full-length transcript protocol exhibit gene length bias akin to bulk RNA-seq data. Specifically shorter genes tend to have lower counts and a higher rate of dropout. In contrast, protocols that include UMIs do not generally exhibit gene length bias. In addition, UMI protocols reveal that shorter genes are as highly expressed as longer genes, and dropout is mostly uniform across genes of varying length. These effects mean that different protocols have the ability to detect a different subset of genes with shorter genes detected more readily using UMI protocols and longer genes detected by full-length protocols.

## Results

### Gene length bias is apparent in scRNA-Seq in non-UMI based protocols

Initially, we analysed three different datasets generated using full-length transcript protocols: mouse embryonic stem cells (Kolodziejczyk et al. 2015), human primordial germ cells (Guo et al. 2015) and human brain whole organoids (Camp et al. 2015). For a full list of the datasets analysed see Supplementary Table 1. Quality control of the single cells was performed and problematic cells filtered out (see ‘Methods’), leaving 530 mouse embryonic stem cells, 226 human primordial germ cells and 494 human brain organoid cells. For each gene, the average log-counts, normalised for sequencing depth, and the proportion of zeroes across the cells (i.e. the dropout rate per gene) were calculated (see ‘Methods’). Gene-wise abundances were estimated for all datasets by dividing the gene-level counts by gene length to obtain Reads Per Kilobase per Million (RPKM). In order to assess gene length bias, genes were assigned to 10 bins based on gene length such that each bin had a roughly 1000 genes. The results are summarised in the boxplots in Figure 1.

**Figure 1:**
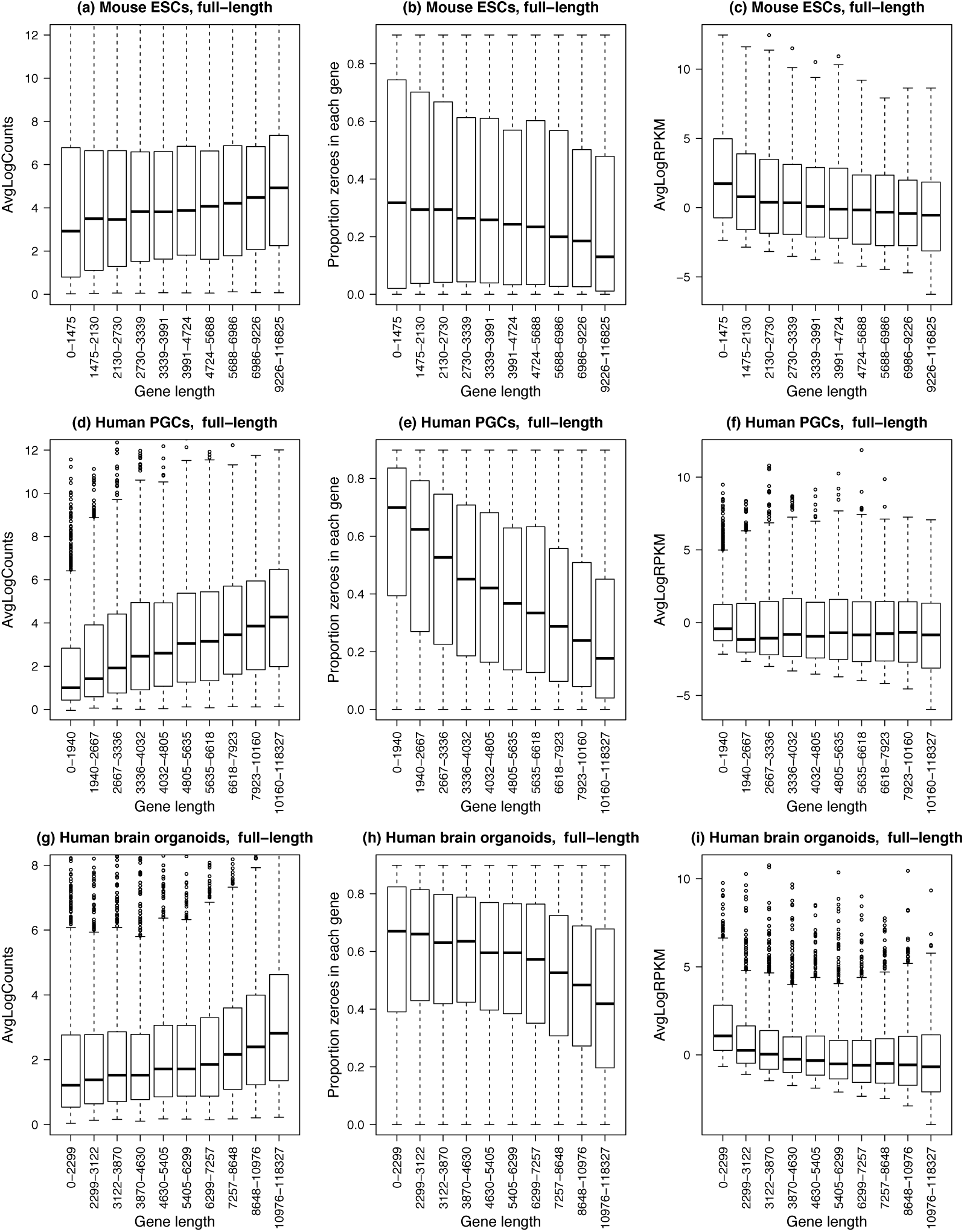
Gene length bias is present in non-UMI protocols. Three different datasets were analysed: (a-c) mouse embryonic stem cells, n=530 (Kolodziejczyk et al. 2015), (d-f) human primordial germ cells, n=226 (Guo et al. 2015), and (g-i) human brain whole organoids, n=494 (Camp et al. 2015). For all plots (a-i), the x-axis shows 10 gene length bins all containing roughly equal numbers of genes. The left panel shows gene-wise average log counts, the middle panel shows proportion of zeroes in each gene (dropout rate per gene), and the right panel shows average log counts corrected for gene length (RPKM).

For all three full-length protocol datasets, shorter genes have lower count level expression proportions compared to longer genes, with a clear trend of increasing log-counts as gene length increases (Figure 1a, d, g). This was accompanied by a decreasing trend in the dropout rate per gene as gene length increased, highlighting the fact that shorter genes are more difficult to detect using full length protocols (Figure 1b, e, h). These trends are stronger for the human PGCs and human brain organoid datasets, while not as severe for the mouse ESCs. Calculating transcript abundance by dividing gene-level counts by gene length mostly removes the gene length bias for the human PGCs and brain organoid datasets (Figure 1f, i), however for the mouse ESCs calculating RPKMs appears to induce a trend with gene length such that shorter genes appear more highly expressed relative to the longer genes (Figure 1c).

### UMI-based protocols do not suffer from gene length bias

We hypothesised that because UMI protocols tag each transcript molecule separately we would not see a similar gene length bias in these protocols. In order to assess gene length bias in scRNA-Seq datasets with included UMIs, we analysed three different datasets: mouse embryonic stem cells generated using a CEL-Seq protocol (Grün et al. 2014; Hashimshony et al. 2012), human induced pluripotent stem cells generated using a modified SMARTer protocol with the Fluidigm C1 system (Tung et al. 2016) and human leukemia cell line K562 cells using the CEL-Seq protocol with InDrop (Klein et al. 2015). After quality control and filtering of problematic cells, 127 single cells remained for the mouse embryonic stem cells, 671 for human induced pluripotent stem cells and 219 human K562 cells (supplementary table 1).

We found that for the human iPSCs and human K562 datasets, the average log-counts were fairly uniform across the 10 gene length bins, and for the mouse ESCs, the shorter genes appear to be more highly expressed than the longer genes (Figure 2a, d, g). Comparing medians, the dropout rate per gene is slightly lower for shorter genes in the mouse ESCs, while for the human iPSCs and K562 cells, the dropout is fairly uniform across the gene length bins, although slightly more variable for the shortest genes (Figure 2b, e, h). However, calculating RPKMs by dividing by gene length induces a clear trend with gene length where shorter genes appear to be more highly expressed relative to longer genes, with the median log RPKM decreasing with increasing gene length (Figure 2c, f, i).

**Figure 2:**
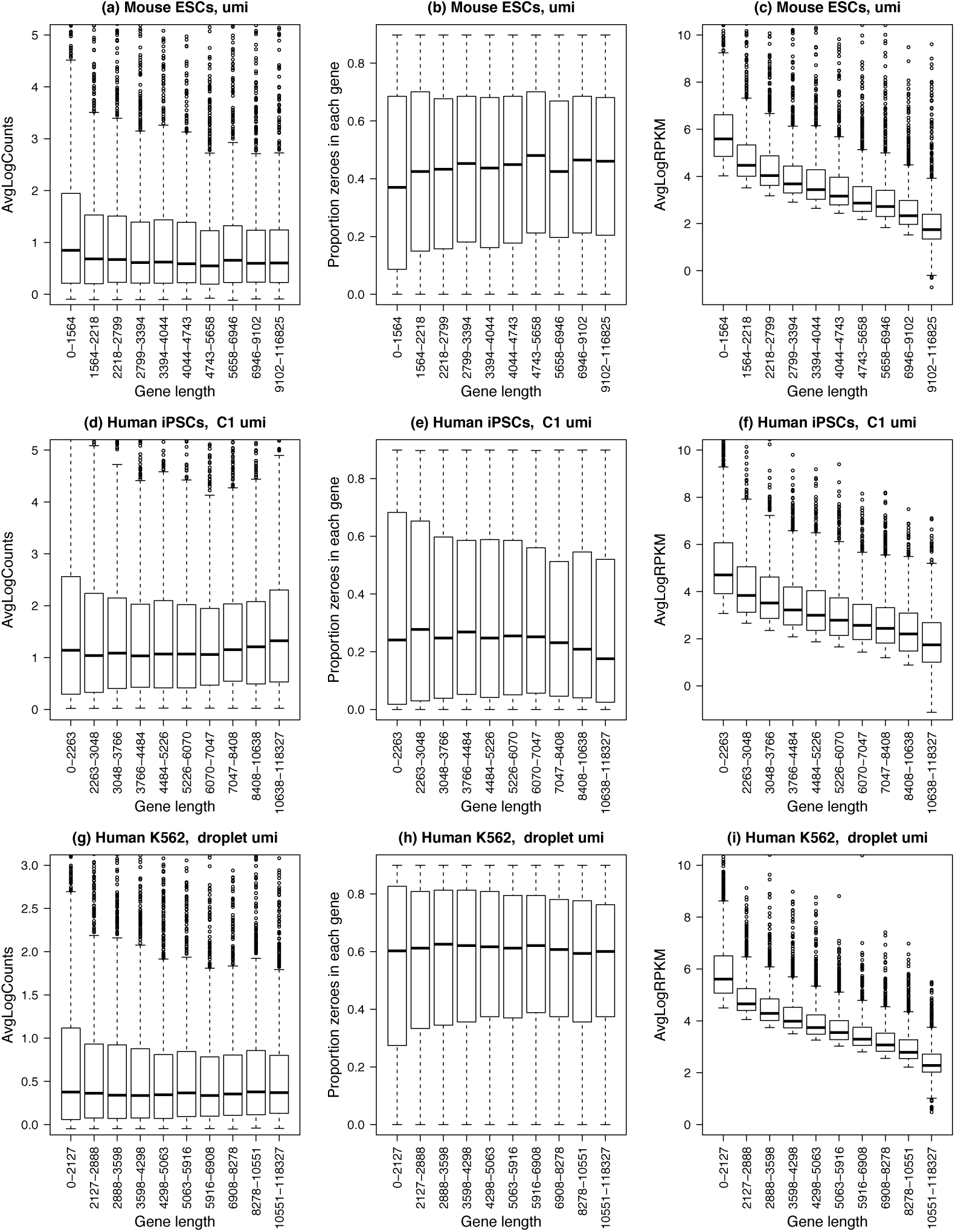
Gene length bias is absent in UMI-based protocols. Three different datasets were analysed: (a-c) mouse embryonic stem cells, n=127 (Grün et al. 2014), (d-f) human induced pluripotent stem cells n=671 (Tung et al. 2016) and (g-i) human leukemia cell line K562 cells, n=219 (Klein et al. 2015). For all plots (a-i), the x-axis shows 10 gene length bins all containing roughly equal numbers of genes. The left panel shows gene-wise average log counts, the middle panel shows proportion of zeroes in each gene (dropout rate per gene), and the right panel shows average log expression corrected for gene length.

### Comparing gene length bias between different mouse embryonic stem cell datasets

To ensure the gene length bias is not due to specific biology of the different cell types, we analysed four different mouse embryonic stem cell datasets generated using both UMI and full-length transcript protocols (Buettner et al. 2015; Ziegenhain et al. 2016; Grün et al. 2014; Kolodziejczyk et al. 2015). When we combined all four datasets together (see ‘Methods’) and performed principal components analysis, we noted that the cells clustered by dataset, with the UMI datasets on the left and full-length datasets on the right of the plot (Figure 3a). Interestingly, in principal components two and three, we saw some biological structure in the datasets emerging, with cells grown in different media clustering together (Figure 3b). In particular, three different datasets (two full-length, one UMI), grown in standard media with 2i inhibitors all cluster together on the left of the plot. This shows great promise for obtaining biologically interesting results from combining multiple datasets generated in different labs using different technology.

**Figure 3:**
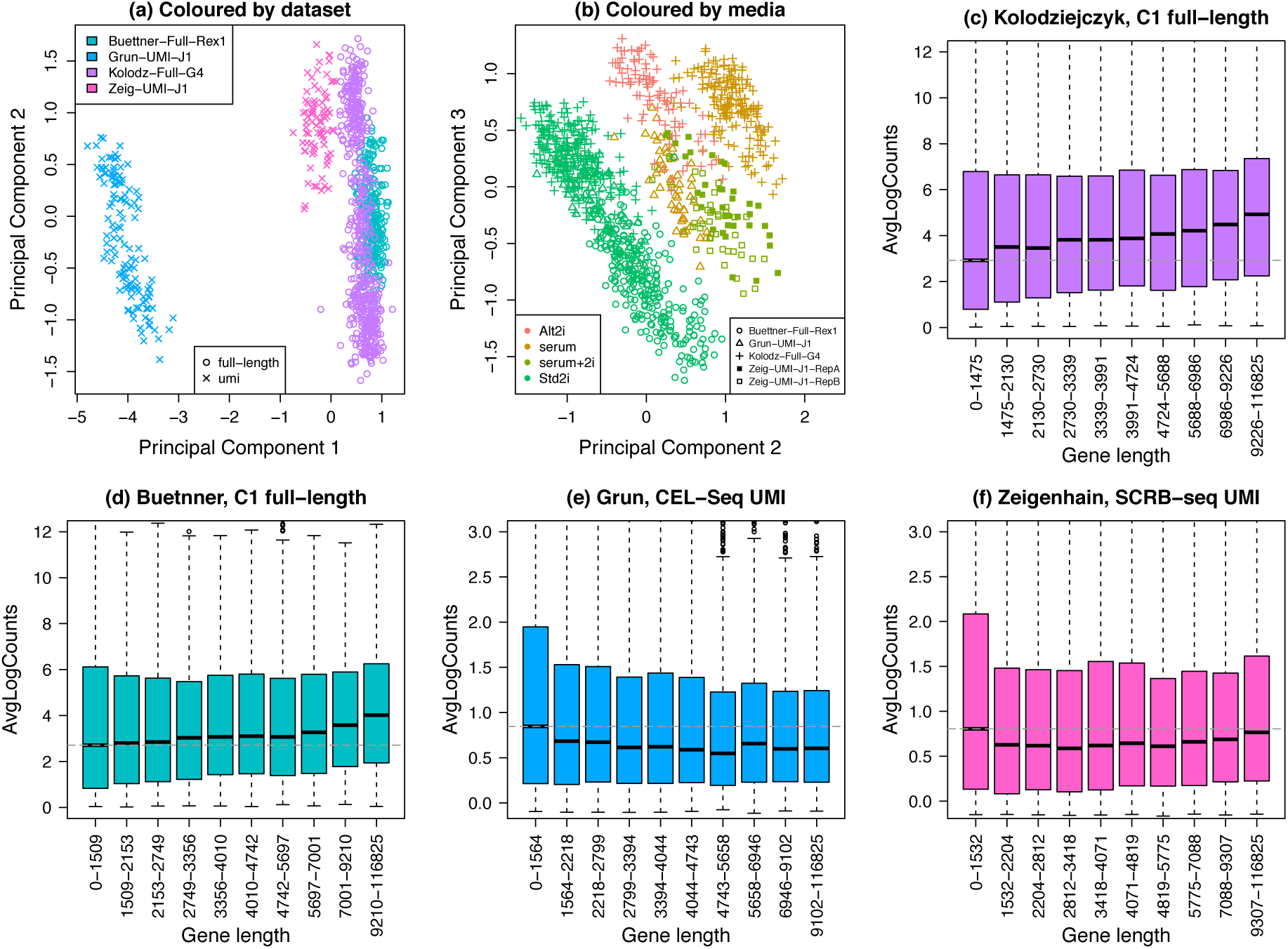
Combining four mouse embryonic stem cell datasets. Four different mouse embryonic stem cell datasets were combined, two full-length transcript (Buettner et al. 2015; Kolodziejczyk et al. 2015) and two UMI datasets (Grün et al. 2014; Ziegenhain et al. 2016). (a) Principal component analysis plot (coloured by dataset) shows the major source of variation between the cells is the dataset, with the UMI datasets on the left and the full-length datasets on the right. (b) Examining principal components two and three reveals that the next major source of variation in the data is the media in which cells are grown. In particular three datasets (two full-length and one UMI) which have cells grown in standard media with 2i inhibitors all cluster together on the left. J1, Rex1 and G4 refer to the mESC cell line. The Ziegenhain dataset has single cells profiled in two batches. (c-d) Gene length bias is present in full-length mESC datasets; dotted grey line is the median log-count in the first gene length bin. (e-f) Gene length bias is absent in UMI mESC datasets; dotted grey line is the median log-count in the first gene length bin.

In terms of the gene length bias across the multiple datasets, it is clear that data generated from full length protocols exhibit gene length bias, with shorter genes having lower average log-counts compared to longer genes (Figure 3c, d). This is not as pronounced compared to other full-length datasets (Figure 1d, g), however compared to the UMI mESC datasets it is quite noticeable. For the UMI datasets, the gene length bias is mostly uniform across the gene length bins, however the shortest genes in the first bin appear to have slightly higher average log-counts and are more variable compared to the longer genes (Figure 3e, f).

### Detection differences in UMI and full-length mESC datasets

In order to investigate whether choice of protocol impacts which genes are detected, we compared genes detected in both UMI mESC datasets to genes detected in both full-length mESC datasets. We found 13434 genes were detected in at least one of the four datasets. Across both UMI datasets, 8866 genes were detected with counts in at least 10% of the cells for each dataset. For the full-length datasets, 11328 genes were detected using the same criteria. The full-length datasets had much greater sequencing depth (median ~ 3million reads, Supplementary Table 1) and more cells compared to the UMI datasets (median ~33,000 reads, Supplementary Table 1), hence it is unsurprising that more genes are detected across both full-length datasets. However, there were 188 genes detected in the UMI datasets that were not detected in the full-length datasets (Figure 4a). The genes unique to the UMI datasets tended to be shorter compared to the gene lengths of the 2644 genes uniquely detected in the full-length datasets (Figure 4b, p-value=0.000297, Wilcoxon Rank Sum Test). The genes uniquely detected in either the full-length or UMI datasets tended to be lowly expressed, hence more difficult to detect in general.

**Figure 4:**
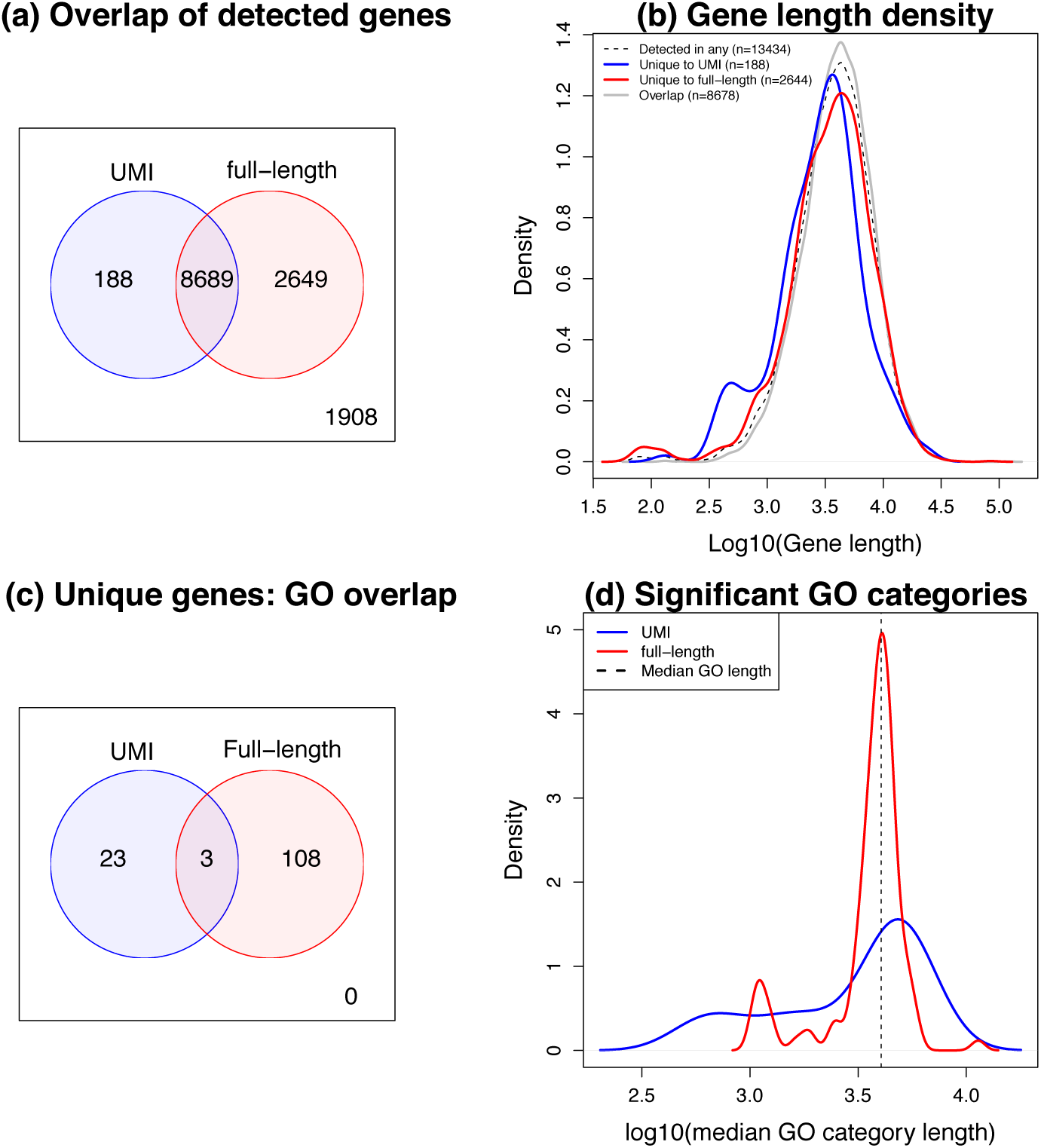
Detection differences in UMI and full-length mESC datasets. (a) A Venn diagram comparing the number of genes detected in two UMI mESC datasets with the number detected in the two full-length datasets. We find that while the majority of genes are detected in all datasets (n=8689), there are genes that are uniquely detected when using either a full-length or UMI protocol. (b) Density plots of gene length for the subsets of genes corresponding to the vennDiagram in (a). The uniquely detected genes for the UMI datasets (blue line) tend to be shorter than the uniquely detected genes in the full-length datasets (red line) p=0.000297. (c) A Venn diagram showing the number of enriched GO categories in the 188 genes unique to UMIs and the 2649 genes unique to the full-length protocols reveals that these genes interrogate different biology, with only 3 GO categories in common. (d) Density plots of average gene length for each GO category corresponding to the significantly enriched GO categories in (c). We assigned each GO category an average length by calculating the median of the lengths of all genes annotated to each GO category. While there is not a significant shift in location in the density plots we noted a much greater spread of median length in the enriched GO categories for the uniquely detected UMI genes, largely driven by the presence of GO categories that tend to have very short genes.

Comparing differential expression between two media (2i inhibitors versus serum) in one UMI dataset (Grün et al. 2014), revealed that 31% (59/188) of the uniquely detected genes were defined as significantly differentially expressed (total differentially expressed = 1641/9962, 16%). For a similar comparison in a full length dataset (2i inhibitors versus serum, (Kolodziejczyk et al. 2015)), 20% (531/2644) of the uniquely detected genes in full –length datasets were significantly differentially expressed (total differentially expressed = 1653/12395, 13%). This highlights that protocol choice may impact ability to detect differential expression of some genes.

Examining which GO terms are over-represented for the 188 genes unique to the UMI dataset revealed that categories such as neural crest cell migration, negative regulation of megakaryocyte differentiation and stem cell development were among the 26 statistically significantly enriched categories (Supplementary Table 2). There were 4/26 GO categories with extremely short average gene length (<1000, median gene length across all GO categories = 4039), with the top two GO categories, “nucleosome” and “DNA packaging complex”, having median gene length in GO categories = 614, 706. However there were also statistically significant categories that are comprised of longer genes than average (13/26 categories with median length > 4039), indicating that pathways enriched for the unique UMI genes were not heavily biased towards categories only containing short genes.

For the full-length datasets, the GO categories that were significantly enriched (n=111) were different to those pathways enriched for the unique UMI genes, with only 3 GO categories overlapping (Figure 4c, Supplementary Table 3). GO categories such as those involved in plasma membrane, cell signalling, ion and cation channel activity were over-represented for the 2649 unique genes. While there were no significantly enriched GO categories that had extremely small average gene length (<1000), 14% (16/111) had median gene length < 2632 (the 5^th^ percentile of median gene length across the GO categories). There was one statistically significant GO category with extremely large average gene length (> 10,000). Although there was no significant shift in median gene length of GO categories between the UMI and full-length GO categories, we noted that the variation in median GO length for the uniquely detected UMI genes was 3.5 times greater than for the uniquely detected full-length genes, largely driven by prevalence of very small sets (Figure 4d, p-value = 5.6×10^−6^, F-test).

## Discussion

While single cell RNA-sequencing technology is advancing at a rapid rate and novel discoveries are being made, the datasets being generated have many technical biases. Here, we have investigated the role that gene length plays in protocols that include UMIs as well as full-length transcript protocols. Unsurprisingly, we find that for full-length protocols, genes that tend to be shorter have lower counts and a higher rate of dropout, while UMI based protocols have a more even distribution of dropout across genes of varying length. In addition, a UMI protocol is more likely to detect lowly expressed genes that are shorter compared to a full-length protocol, where lowly expressed genes that are longer are easier to detect (Supplementary Figure 1). Of course UMI protocols are unable to provide information on transcript structure such as which isoforms are expressed in a sample and only provide overall gene level expression measures. Since UMI counts are already molecule counts, expression levels should be expressed as normalised counts (e.g. counts per million) rather than dividing by gene length to obtain RPKMs, as this latter measure will artificially inflate the expression estimates of shorter genes relative to longer genes.

While datasets generated using a UMI based protocol tend to have much lower sequencing depths, and hence lower counts, we found that in mESCs we were still able to detect uniquely expressed genes in the UMI datasets that were not detected in full-length datasets. However, a larger set of genes were detected in the mESC full-length datasets. Performing GO analysis on genes uniquely detected by each protocol revealed that they interrogate different biology, and hence the choice of protocol may affect which pathways can be studied. In particular, the genes unique to either the UMI or the full-length datasets appeared to be biologically relevant, as a subset were found to be significantly differentially expressed when comparing cells grown in two different media.

We combined four different datasets generated from mESCs that had strikingly different sequencing depths and protocols. Despite these differences, we found that we were able to recover biologically relevant structure. In particular, three different datasets (two full-length, one UMI), grown in standard media with 2i inhibitors all cluster together when examining higher principal components. Although promising, the greatest source of variation between the cells was the dataset they belonged to highlighting the known issues with large batch effects in scRNA-seq (Tung et al. 2016; Hicks et al. 2015). Hence, analysis methods including data cleaning and normalisation are crucial when combining datasets in order to extract biologically meaningful relationships.

## Methods

### Processing

We processed three datasets through our pipeline developed for full-length data: mouse embryonic stem cells (Kolodziejczyk et al. 2015), human cerebral organoid cells (Camp et al. 2015), and mouse embryonic stem cells (Buettner et al. 2015). The quality of the raw sequencing reads was examined using FastQC v0.11.4 and they were checked for contamination by aligning a sample of reads to multiple reference genomes using FastQ Screen v0.6.4. Reads were aligned to the appropriate reference genome (mm10 for mouse and hg38 for human), using STAR v2.5.2a (Dobin et al. 2013) and reads overlapping genes in an appropriate annotation (Gencode M9 for mouse, Gencode V22 for human) were counted using featureCounts v1.5.0-p3 (Liao et al. 2014). This pipeline was constructed in Bpipe v0.9.9.3 (Sadedin et al. 2012) and a report summarising the steps produced using MultiQC v0.8 (Ewels et al. 2016).

### Gene filtering

Our gene filtering strategy was identical between datasets. Genes that had more than 90% zeroes across all cells, as well as ribosomal and mitochondrial genes were filtered out. Genes that could not be annotated with an Entrez Gene ID were also removed in order to retain a set of well curated genes. Gene length information was taken as the sum of the exon lengths as outputted by the featureCounts software for mm10 GENCODE VM4 annotation for all mouse datasets, and for all human datasets, we used the sum of exon lengths as outputted by featureCounts for the hg38 GENCODE V22 annotation. Genes that could not be annotated with gene length information were filtered out. We found that using these criteria helped reduce some of the variability in the datasets.

### Datasets

The details of all datasets analysed are included in Supplementary Table 1.

#### Mouse embryonic stem cells, Kolodziejczyk et al. 2015, full-length

We downloaded the raw data from the ArrayExpress database (http://www.ebi.ac.uk/arrayexpress) under accession number E-MTAB-2600 and ran our full-length processing pipeline using the mm10 mouse genome to produce a counts matrix. We performed quality control on the cells and removed cells that had a dropout rate of greater than 80% and a library size of fewer than half a million. We calculated the proportion of sequencing reads taken up by the ERCC spike-ins and discarded three plates that had proportions of ERCC spike-ins that appeared excessive compared to the remaining plates. We performed gene filtering as described above. After cell and gene filtering, we were left with 530 cells and 12395 genes for further analysis.

#### Human primordial germ cells, Guo et al. 2015, full-length

We downloaded the processed data from the Conquer website (http://imlspenticton.uzh.ch:3838/conquer/). The data had been pseudo-aligned to the latest human reference genome, hg38, using the Salmon software tool. The data is also available under the GEO accession number GSE63818. There did not appear to be any spike-in controls for this dataset, hence filtering was performed on the dropout rate and total sequencing depth for each cell. Cells with more than 85% dropout and fewer than half a million sequencing reads were filtered out. After cell and gene filtering, there were 226 cells and 15837 genes for further analysis.

#### Human cerebral organoid cells, Camp et al. 2015, full-length

We downloaded the data from SRA (SRP066834) and ran our full-length processing pipeline to produce a counts matrix, using the hg38 human genome. We removed cells that had greater than 90% dropout, library size smaller than half a million as well as cells that had more than 20% of the sequencing taken up by ERCC controls. After cell and gene filtering, we had 494 cells and 11325 genes for further analysis.

#### Mouse embryonic stem cells, Grün et al. 2014, UMI

We downloaded the processed data from GEO under accession number GSE54695. The data was aligned to the mouse genome (mm10) using BWA and transcript number estimated from UMI counts by the authors. We removed cells that had > 80% dropout, library size smaller than 10000, as well as cells that had more than 5% of the sequencing taken up by ERCC controls. After cell and gene filtering, there were 127 cells and 9962 genes for further analysis.

#### Human induced pluripotent stem cells, Tung et al. 2016, UMI

We downloaded the processed molecule counts and sample information from the authors’ github repository (https://github.com/jdblischak/singleCellSeq). The data was aligned to the human genome (hg19) using the Subjunc aligner. The data is also available under GEO accession GSE77288. We removed cells that had > 70% dropout, fewer than 30000 sequencing reads per cell, as well as cells that had more than 3% of the sequencing taken up by ERCC spike-ins. After cell and gene filtering, we had 671 cells and 11971 genes for further analysis.

#### Human K562 cells (lymphoblastoma culture), Klein et al. 2015, UMI

The processed molecule count data was downloaded from GEO under accession GSM1599500. The data was aligned to the human genome (hg19) using Bowtie. Cells that had > 85% dropout, fewer than 10000 total sequencing reads, or an ERCC library size to total library size ratio > 0.01 were filtered out. After cell and gene filtering, we had 219 cells and 13418 genes for further analysis.

#### Mouse embryonic stem cells, Zeigenhain et al. 2016, UMI

We downloaded the molecule counts from GEO under accession GSE75790. The SCRB-Seq protocol was used to generate the libraries, and the data had been processed by the authors through a dropseq pipeline, which included alignment to the mouse genome (mm10) using STAR. The cells all appeared good quality hence cell filtering wasn’t necessary. After gene filtering, we had 84 cells and 10519 genes for further analysis.

#### Mouse embryonic stem cells, Buettner et al. 2015, full-length

We downloaded the data from the European Nucleotide Archive,http://www.ebi.ac.uk/ena/data/view/PRJEB6989, and ran the data through our full-length pipeline, mapping to the mouse genome (mm10) to produce a counts matrix. We filtered out cells with > 85% dropout and sequencing depth less than a million. After cell and gene filtering, we had 271 cells and 11700 genes for further analysis.

### Combining mouse embryonic stem cell datasets

We combined four different mouse embryonic stem cell datasets using the following approach. We performed gene and cell filtering on each dataset independently, and combined the datasets by taking the genes commonly detected across all four datasets (8678 genes, 1012 cells, each gene is detected in at least 10% of the cells for each dataset). This strategy ensured that the genes were detected in all four datasets, and hence larger datasets did not dominate gene filtering. It also ensured that the larger datasets did not dominate the principal components analysis.

### Statistical analysis

All statistical analysis was performed in R-3.3.1, using the limma (Ritchie et al. 2015), edgeR (Robinson et al. 2010), scran (Lun et al. 2016) and scater (McCarthy et al. 2016) Bioconductor packages (Gentleman et al. 2004). The UMI dataset was normalised using scran prior to differential expression analysis, as it clearly showed composition bias. Differential expression analysis in the mESCs was performed using edgeR, specifying a log-fold-change cut-off of 1 for the full-length dataset, and 0.5 for the UMI dataset. GO analysis was performed with hypergeometric tests using the goana function in limma. All scripts for analysing the datasets are available on the Oshlack lab github page (https://github.com/Oshlack/GeneLengthBias-scRNASeq).

## Acknowledgements

Luke Zappia is supported through an Australian Government Research Training Program Scholarship. Alicia Oshlack is supported through an NHMRC Career Development Fellowship APP1126157.

## Supplementary Figures

**Supplementary Figure 1: Average log counts for detected genes in UMI and full-length transcript protocols.** The average log counts tend to be much lower for UMI dataset compared to the full-length datasets. The genes uniquely detected for each protocol tend to be lowly expressed, hence more difficult to detect.

## Supplementary Tables

**Supplementary Table 1:** Details of the datasets analysed in the paper.

**Supplementary Table 2:** Enrichment of GO categories for the 188 genes uniquely detected in the UMI mESC datasets.

**Supplementary Table 3:** Enrichment of GO categories for the 2649 genes uniquely detected in the full-length mESC datasets.

